## Supplemental Information for "SPEARS: Standard Performance Evaluation of Ancestral Reconstruction through Simulation"

Heather Manching and Randall J. Wisser*

Department of Plant & Soil Sciences, University of Delaware, Newark, DE, USA

### Tables

**Table S1. SPEARS Results for Low and High Marker Density.**

| Markers | AAA[a] | GAA[b] | SER[c] | CCC[d] |
|---|---|---|---|---|
| $n$ = 23,584 | 97.3% ± 0.6% | 99.6% ± 0.1% | 0.0098 ± 0.0018 | $r$ = 0.89, $p$ < 2.2 e-16 |
| $n$ = 47,074 | 97.0% ± 0.6% | 99.3% ± 0.1% | 0.0090 ± 0.0018 | $r$ = 0.89, $p$ < 2.2 e-16 |

[a]Ancestral assignment accuracy plus-minus 1 standard deviation.
[b]Genotype assignment accuracy plus-minus 1 standard deviation.
[c]Switch error rate plus-minus 1 standard deviation.
[d]Correlation between crossover counts.

### Figures

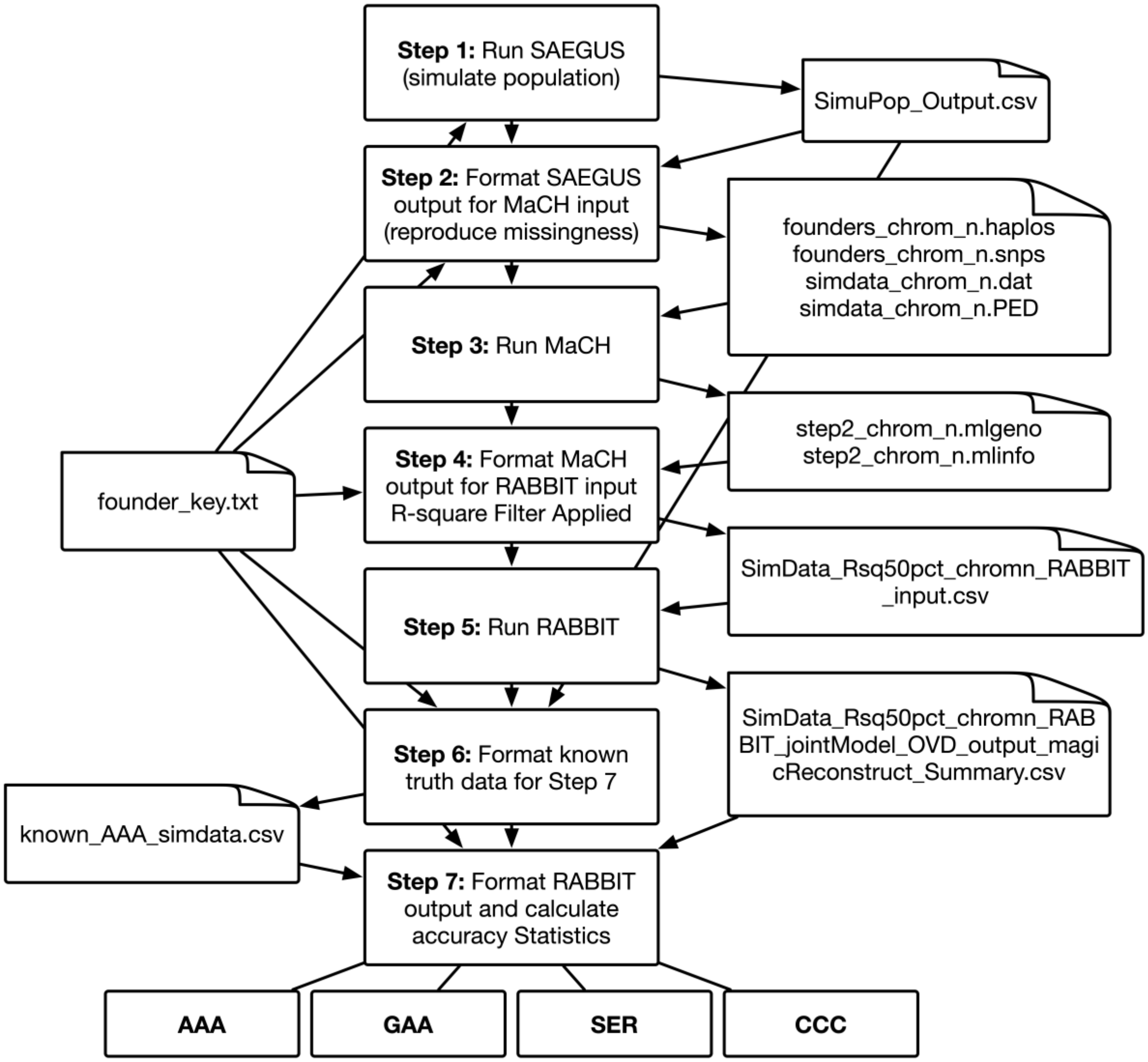

**Figure S1. SPEARS Pipeline.** The figure shows the SPEARS pipeline, which begins with the creation of a simulated population (*N* virtual genotypes) using SAEGUS based on a user-provided genetic map and parental genotype data. Simulated output is formatted for use in MaCH and RABBIT based on parental genotype data. Finally, output from RABBIT is compared to original simulated data to calculate accuracy statistics: AAA, GAA, SER, and CCC (see methods in main text).

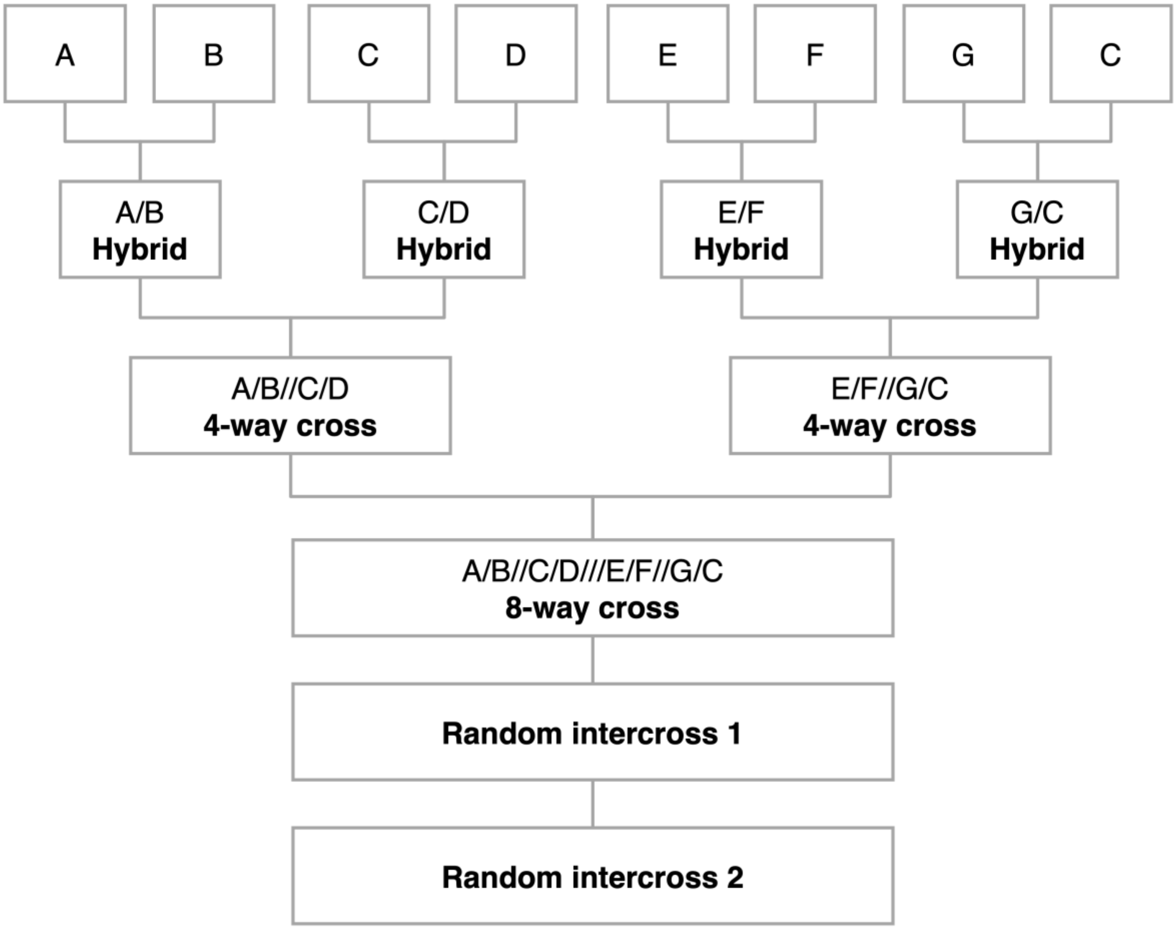

**Figure S2. Multi-Parent Pedigree used to Demonstrate SPEARS.** As proof-of-concept for the study, mimicking the pedigree of a real population of maize, multiple inbred line parents were used to create virtual genotypes (parent C was used twice). Each generation was created by randomly intermating according to the pedigree. One-thousand progeny from the last intercross were used for SPEARS. See the code for this example at github (https://github.com/maizeatlas/spears/blob/master/1_SAEGUS.py) for detailed parameters used for the simulation in SAEGUS.

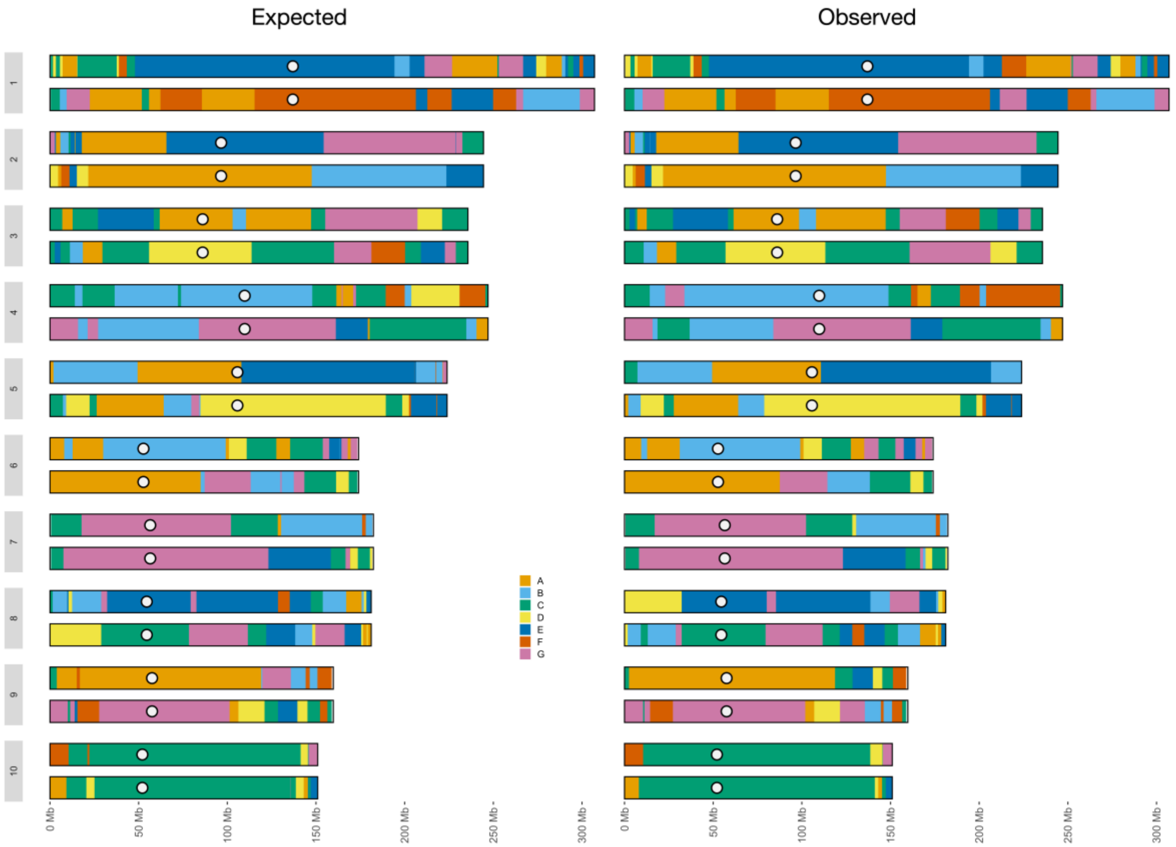

**Figure S3. Ancestral Haplotype Maps from SPEARS.** The expected (simulated) and observed (RABBIT-inferred) haplotype map for all chromosomes in a single individual is shown. Colors correspond to the seven founding parents of the admixed population (Figure S2). The centromere is represented as a white circle.

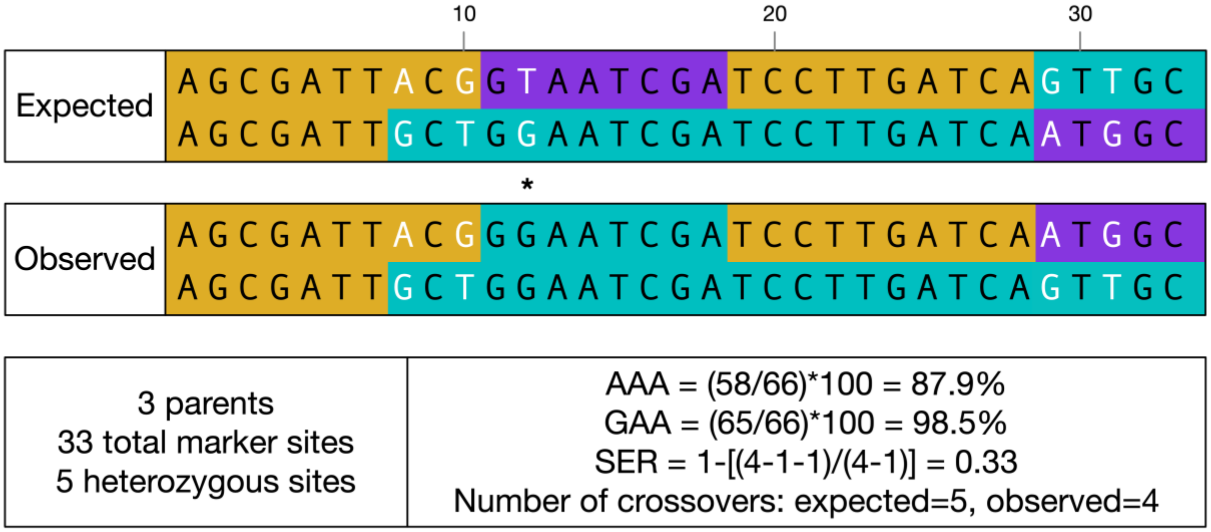

**Figure S4. SPEARS Metrics.** The figure portrays diplotypes for a segment of the genome and describes how each summary statistic is calculated. Each color represents a distinct parent-of-origin. Heterozygous genotypes are shown in white text. The top diplotype represents expected data (output from simulation) and the bottom diplotype represents observed data (e.g., output from RABBIT). AAA and GAA are phase independent and are calculated as the proportion of marker sites ($n$ = 33) that have the correct parent assignment or genotype assignment, respectively. Sites with genotype errors are marked with an asterisk. SER is calculated as (1-($n$-$sw$-1)/($n$-1)), where $n$ is the total number of heterozygous sites (four in this case) and $sw$ is the number of switches required to obtain the correct phase; heterozygous sites with genotype errors are excluded from SER calculation. Finally, CCC is calculated as the correlation of crossover counts (shown here) for all samples in the simulated output.

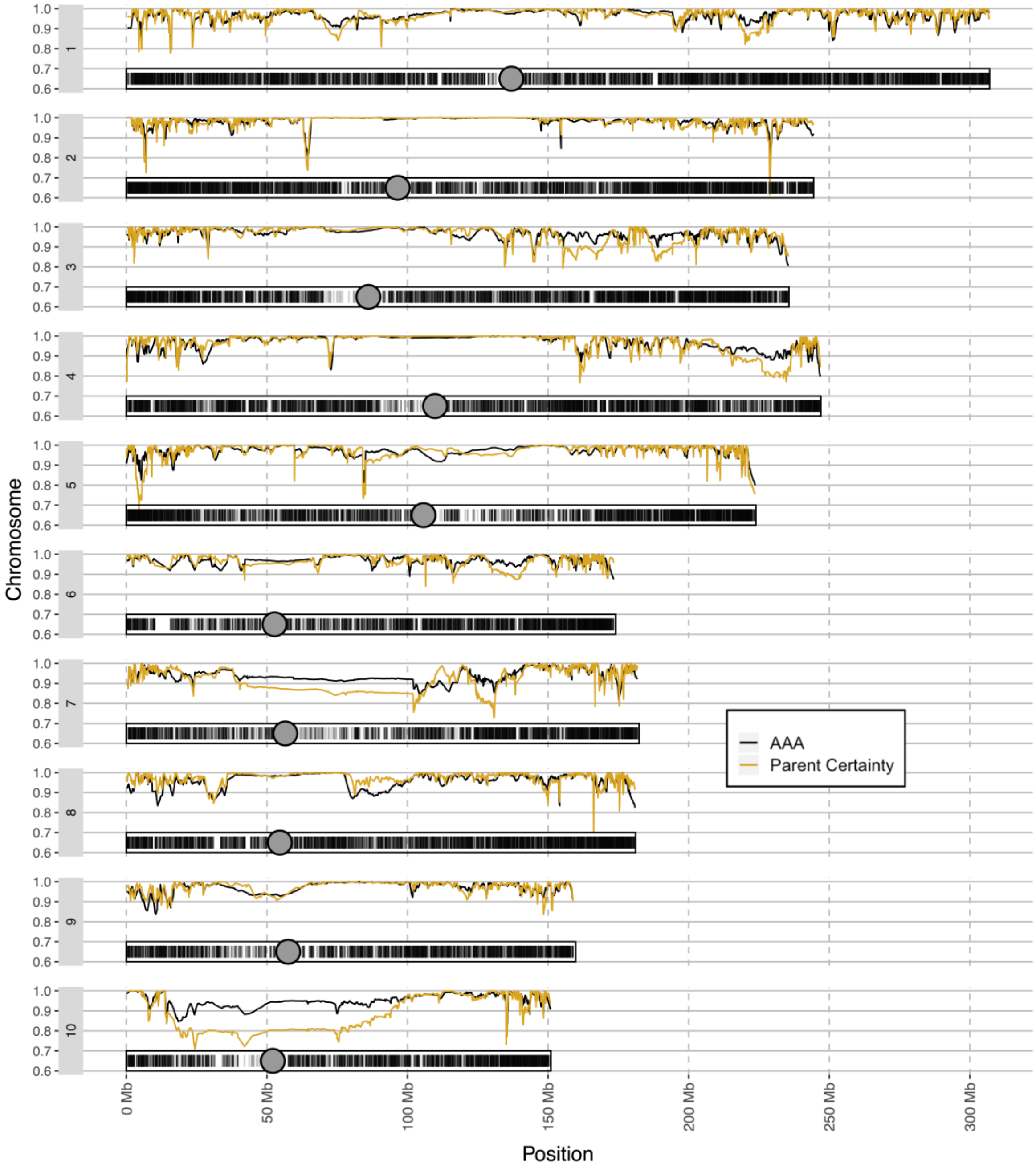

**Figure S5. Genome-wide Ancestral Assignment Accuracy.** AAA (black), parent certainty (gold), marker density (rug along x-axis), and centromere position (grey dot with black outline) were plotted for each of the ten chromosomes of maize. Parent certainty is the difference in parent probabilities for the two most likely parents at each marker, where a higher value indicates more certainty in assignment of the parent (i.e., a larger difference between the two most likely parents).

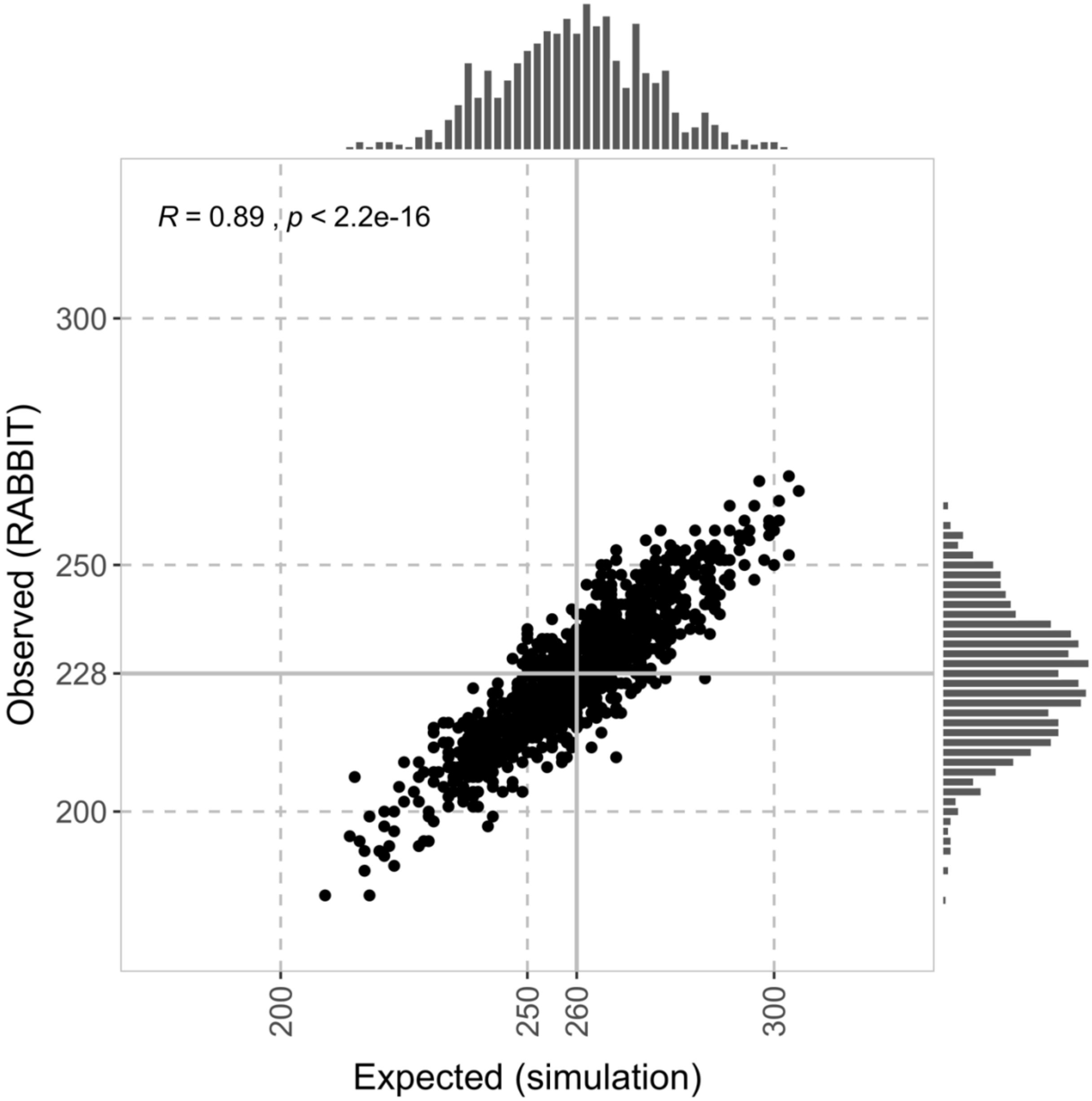

**Figure S6. Correlation and Distributions of Crossover Counts** Each point is the number of crossovers for a given individuals observed versus expected value. Solid grey lines indicate the average number of crossovers among individuals. Marginal histograms show the distributions in crossover counts.

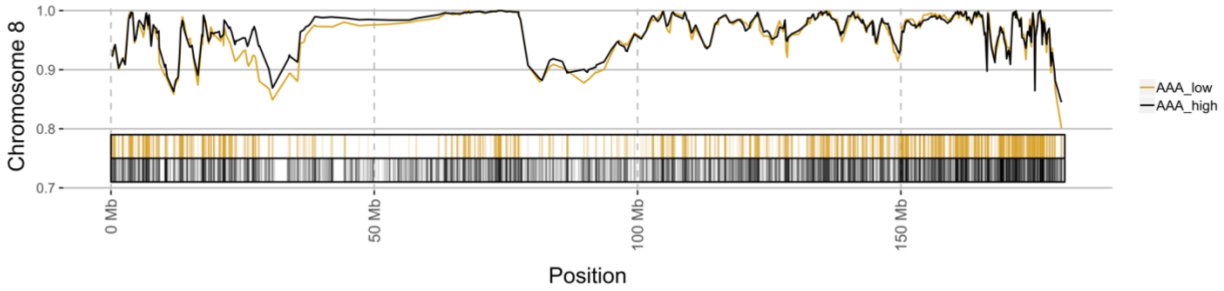

**Figure S7. Ancestral Assignment Accuracy for Low and High Marker Densities.** AAA is shown for SPEARS using *n* = 23,584 markers (gold) and *n* = 47,074 markers (black) along with the corresponding marker densities (rugs along x-axis) for chromosome 8. The region between 25 Mb and 50 Mb demonstrates an increase in accuracy in relation to marker density. The region between 75 Mb and 100 Mb demonstrates minimal change in accuracy despite an increase in marker density and is likely due to IBS/IBD.
