## Supplementary material for "SPEARS: Standard Performance Evaluation of Ancestral Reconstruction through Simulation": LaTex files from Overleaf: Article.tex

\documentclass[times, twoside, watermark]{zHenriquesLab-StyleBioRxiv}
\usepackage{blindtext}

% Please give the surname of the lead author for the running footer
\leadauthor{Manching} 

\begin{document}

\title{SPEARS: Standard Performance Evaluation of Ancestral Reconstruction through Simulation}
\shorttitle{Appraising Ancestral Haplotype Reconstruction}

% Use letters for affiliations, numbers to show equal authorship (if applicable) and to indicate the corresponding author
\author[1,]{H. Manching}
\author[1,\Letter]{R. J. Wisser}

\affil[1]{Department of Plant \& Soil Sciences, University of Delaware, Newark, DE, 19716 USA}

\maketitle

%TC:break Abstract
%the command above serves to have a word count for the abstract
\begin{abstract}
%\blindtext
\textbf{Motivation:} Ancestral haplotype maps provide useful information about genomic variation and biological processes. Reconstructing the descendent haplotype structure of homologous chromosomes, particularly for large numbers of individuals, can help with characterizing the recombination landscape, elucidating genotype-to-phenotype relationships, improving genomic predictions and more. Inferring haplotype maps from sparse genotype data is an efficient approach to whole-genome haplotyping, but this is a non-trivial problem. A standardized approach is needed to validate whether haplotype reconstruction software, conceived population designs and existing data for a given population provides accurate haplotype information for further inference.\\
\newline
\textbf{Results:} We introduce SPEARS, a pipeline for whole simulation-based appraisal of genome-wide ancestral haplotype inference. The pipeline generates virtual genotypes (truth data) with real-world missing data structure. It then proceeds to mimic analysis in practice, capturing sources of error due to imputation and reconstruction of ancestral haplotypes. Standard metrics allow researchers to assess which features of haplotype structure or regions of the genome are sufficiently accurate for analysis and reporting. Haplotype maps for 1,000 outcross progeny from a multi-parent population of maize is used to demonstrate SPEARS.\\
\newline
\textbf{Availability:}  \href{https://github.com/maizeatlas/spears}{https://github.com/maizeatlas/spears}
\end {abstract}
%TC:break main
%the command above serves to have a word count for the abstract

\begin{keywords}
Haplotype reconstruction | Meiotic recombination | Genotype imputation | Genome simulator | RABBIT software
\end{keywords}

\begin{corrauthor}
%\texttt{rjw{@}udel.edu}
rjw\at udel.edu
\end{corrauthor}

\section*{Introduction}
The genome is a mosaic of ancestral haplotypes that capture the evolutionary or breeding history of an individual. Reconstructing ancestral blocks is important for imputing untyped regions of the genome \citep{davies2016rapid}, mapping quantitative trait loci \citep{mott2000method}, investigating the recombination landscape \citep{morgan2017structural}, and inferring the evolutionary history and structure of haplotypes \citep{gabriel2002structure,aylor2011genetic}. Accurately inferring ancestral haplotypes is non-trivial, but several approaches for this have been developed: MERLIN \citep{abecasis2002merlin}; HAPPY \citep{mott2000method}; GAIN \citep{liu2010efficient}; DOQTL \citep{gatti2014quantitative}; R/qtl2 \citep{broman2019r}. These tools have been shown to perform well, but are mostly limited to specific population types or breeding schemes and can be computationally intensive with complex pedigrees or large numbers of markers.
 
RABBIT (Reconstructing Ancestry Blocks BIT by bit) is a flexible tool that uses a Markovian model to reconstruct haplotypes for complex pedigrees involving various mating scenarios \citep{zheng2015reconstruction}. Compared to current tools, it has shown the highest accuracy for inbred lines from multiparent pedigrees (including the mouse CC design [\citealp{churchill2004collaborative}] and \textit{Arabidopsis thaliana} MAGIC design [\citealp{kover2009multiparent}]) \citep{zheng2015reconstruction}. Here, tailored for RABBIT but extensible for other tools, we present a standard performance evaluation of ancestral reconstruction through simulation (SPEARS) pipeline. SPEARS is designed to determine expectations for the accuracy of haplotype reconstruction software and different population designs. As proof-of-concept and a new demonstration of RABBIT, we develop a detailed picture on the variation in accuracy for genome-wide haplotype maps for progeny from a multi-parent population of maize. 

\section*{Materials and methods}
\subsection{Simulation Data}
This study introduces SPEARS (Figure S1) which incorporates SAEGUS \\  (\href{https://github.com/maizeatlas/saegus}{https://github.com/maizeatlas/saegus}) as the genome simulator. Using SAEGUS, we first generated a virtual multi-parent outcross population of $1,000$ progeny (Figure S2). Real genotype data on inbred line parents of the population was used to initiate the simulation ($n=47,078$ markers obtained from both genotyping-by-sequencing (GBS) [\citealp{manching2017phased}] and the MaizeSNP50 BeadChip [\citealp{ganal2011large}]; filtered to remove residual heterozygous sites in the parents for compatibility with MaCH imputation [\citealp{li2010mach}]). We also examined the impact of using fewer markers ($n=23,584$ GBS markers).

In order to represent analysis in practice, the level of missing data per marker observed in a real maize population (not shown) was reproduced for the virtual progeny. That is, at each marker, virtual genotype scores were randomly converted to missing according to the missing data rate observed at the corresponding marker. These missing data were then imputed using MaCH \citep{li2010mach} and filtered to exclude markers with imputation accuracy $<0.5$, resulting in $47,074$ markers (or $23,584$ GBS markers) being retained for RABBIT.

\begin{figure*}[htp]
\renewcommand{\familydefault}{\sfdefault}\normalfont
\centering
\includegraphics[width=\linewidth]{Figure_1.pdf}
\caption{Ancestral Haplotype Maps from SPEARS. The expected (simulated) and observed (RABBIT-inferred) haplotype map for one chromosome in a single individual is shown (for all chromosomes, see Figure S$3$). Colors correspond to the seven founding parents of the admixed population (Figure S2). The centromere is represented as a white circle.}
\label{fig:figure1}
\end{figure*}
%\vspace*{-12pt}

\subsection{Reconstruction of Ancestral Haplotypes}

To build haplotype maps for the simulated population (Figure \ref{fig:figure1}, Figure S3) the joint model of RABBIT was used to assign an optimal Viterbi path using the origViterbiDecoding algorithm \citep{zheng2015reconstruction}. Custom scripts in R were used to reformat output from RABBIT into phased genotypes for SPEARS.

\subsection{Evaluation of Ancestral Reconstruction}

Based on comparing simulated and inferred data per individual, SPEARS uses four metrics to assess distinct features of haplotype maps (Figure S4): (i) Ancestral Assignment Accuracy (AAA); (ii) Genotype Assignment Accuracy (GAA); (iii) Switch-Error Rate (SER); and (iv) Correlation between Crossover Counts (CCC). AAA is calculated as the proportion of markers that have the correct inferred parent assigned on each homologue (given as a percentage). To calculate GAA, genotypes are assigned based on the inferred ancestry and corresponding parental genotype data used as input, from which the proportion of genotype matches are calculated (given as a percentage). To assess phasing accuracy, SER is calculated only among markers with correctly inferred genotype scores (phasing cannot be assessed for markers with genotyping errors) and is defined as: 1-(\textit{n}-1-\textit{sw})/(\textit{n}-1) where \textit{n} is the number of heterozygous sites and \textit{sw} is the number of switches required to obtain the true phase based on known data \citep{stephens2003comparison, marchini2006comparison}. The CCC is calculated as the correlation coefficient for the total number of crossovers across both homologues of all chromosomes per individual. For additional analysis, we also calculated ``parent certainty'' as the difference between the posterior probabilities of the two most likely parents at each marker (obtained from the origPosteriorDecoding algorithm within RABBIT).

\section*{Results}

SPEARS metrics are reported in Table S1. The average per sample AAA was $97.0\%$. Regions with lower accuracy showed decreases in parent certainty alone or in combination with a lower density of markers (Figure S5), indicating that both identity-by-state among parents and marker density, but neither RABBIT nor the Viterbi algorithm per se were main sources of error in reconstructing haplotype maps. Additionally, the average GAA ($99.3\%$) was higher than AAA (Table S1), suggesting shared parental haplotypes were indeed a contributing factor to error. RABBIT performed very well in phasing among markers that had correctly assigned genotypes with an average SER of $0.0090$ across all samples. The CCC was positive and showed high correlation between known and RABBIT-inferred crossover counts (\textit{r} = $0.89$, $p < 2.2 e-16$) (Figure S6). However, the number of crossover counts per sample was downward biased with an average of $260\pm16$ versus $228\pm14$ for known and RABBIT data, respectively.

\section*{Conclusions}

Reconstruction of ancestral haplotypes from genomic data is useful for a number of applications. SPEARS showed that RABBIT produced highly accurate genome-wide haplotype maps for an admixed non-inbred population. Genomic segments with minimal differentiation between the parents resulted in localized reductions in the accuracy of ancestral assignment (Figure S5). This may be overcome with a higher marker density unless parental haplotypes are truly identical-by-descent. For this study, using approximately half the number of markers had minimal influence on the average genome-wide AAA, GAA, CCC, and SER (Table S1); some local regions showed slight reductions in accuracy (Figure S7). This exemplifies how SPEARS estimates of the expectation for different types of error can help guide investigations on haplotype structure. It allows analysis of certain features of haplotype data to be included/excluded in a study based on a corresponding metric, and the expectations can be reported. Moreover, one can assess whether specific chromosomes or regions of the genome, but not others, are sufficiently accurate for downstream analysis. SPEARS, the protocol and suite of scripts, has been made publicly available at \href{https://github.com/maizeatlas/spears}{https://github.com/maizeatlas/spears}.

\begin{acknowledgements}
We thank Dr. Chaozhi Zheng for support in operating RABBIT. \vspace*{-12pt}
\end{acknowledgements}

\section*{Funding}
This work was supported by the Agriculture and Food Research Initiative Competitive Grant (grant no. 2011-67003-30342); and the Agriculture and Food Research Initiative Fellowships Grant Program (grant no. 2018-67011-28052) from the United States Department of Agriculture National Institute of Food and Agriculture.\vspace*{-12pt}

\section*{Bibliography}
\bibliography{spears_citations}

\end{document}
