## Supplementary material for "SPEARS: Standard Performance Evaluation of Ancestral Reconstruction through Simulation": LaTex files from Overleaf: spears_citations.bib

@article{gabriel2002structure,
  title={The structure of haplotype blocks in the human genome},
  author={Gabriel, Stacey B and Schaffner, Stephen F and Nguyen, Huy and Moore, Jamie M and Roy, Jessica and Blumenstiel, Brendan and Higgins, John and DeFelice, Matthew and Lochner, Amy and Faggart, Maura and others},
  journal={Science},
  volume={296},
  number={5576},
  pages={2225--2229},
  year={2002},
  publisher={American Association for the Advancement of Science}
}

@article{gatti2014quantitative,
  title={Quantitative trait locus mapping methods for diversity outbred mice},
  author={Gatti, Daniel M and Svenson, Karen L and Shabalin, Andrey and Wu, Long-Yang and Valdar, William and Simecek, Petr and Goodwin, Neal and Cheng, Riyan and Pomp, Daniel and Palmer, Abraham and others},
  journal={G3: Genes, Genomes, Genetics},
  volume={4},
  number={9},
  pages={1623--1633},
  year={2014},
  publisher={G3: Genes, Genomes, Genetics}
}

@article{kover2009multiparent,
  title={A multiparent advanced generation inter-cross to fine-map quantitative traits in Arabidopsis thaliana},
  author={Kover, Paula X and Valdar, William and Trakalo, Joseph and Scarcelli, Nora and Ehrenreich, Ian M and Purugganan, Michael D and Durrant, Caroline and Mott, Richard},
  journal={PLoS genetics},
  volume={5},
  number={7},
  year={2009},
  publisher={Public Library of Science}
}

@article{li2010mach,
  title={MaCH: using sequence and genotype data to estimate haplotypes and unobserved genotypes},
  author={Li, Yun and Willer, Cristen J and Ding, Jun and Scheet, Paul and Abecasis, Gon{\c{c}}alo R},
  journal={Genetic epidemiology},
  volume={34},
  number={8},
  pages={816--834},
  year={2010},
  publisher={Wiley Online Library}
}

@article{liu2010efficient,
  title={Efficient genome ancestry inference in complex pedigrees with inbreeding},
  author={Liu, Eric Yi and Zhang, Qi and McMillan, Leonard and De Villena, Fernando Pardo-Manuel and Wang, Wei},
  journal={Bioinformatics},
  volume={26},
  number={12},
  pages={i199--i207},
  year={2010},
  publisher={Oxford University Press}
}

@article{manching2017phased,
  title={Phased genotyping-by-sequencing enhances analysis of genetic diversity and reveals divergent copy number variants in maize},
  author={Manching, Heather and Sengupta, Subhajit and Hopper, Keith R and Polson, Shawn W and Ji, Yuan and Wisser, Randall J},
  journal={G3: Genes, Genomes, Genetics},
  volume={7},
  number={7},
  pages={2161--2170},
  year={2017},
  publisher={G3: Genes, Genomes, Genetics}
}

@article{morgan2017structural,
  title={Structural variation shapes the landscape of recombination in mouse},
  author={Morgan, Andrew P and Gatti, Daniel M and Najarian, Maya L and Keane, Thomas M and Galante, Raymond J and Pack, Allan I and Mott, Richard and Churchill, Gary A and de Villena, Fernando Pardo-Manuel},
  journal={Genetics},
  volume={206},
  number={2},
  pages={603--619},
  year={2017},
  publisher={Genetics Soc America}
}

@article{mott2000method,
  title={A method for fine mapping quantitative trait loci in outbred animal stocks},
  author={Mott, Richard and Talbot, Christopher J and Turri, Maria G and Collins, Allan C and Flint, Jonathan},
  journal={Proceedings of the National Academy of Sciences},
  volume={97},
  number={23},
  pages={12649--12654},
  year={2000},
  publisher={National Acad Sciences}
}

@article{stephens2003comparison,
  title={A comparison of bayesian methods for haplotype reconstruction from population genotype data},
  author={Stephens, Matthew and Donnelly, Peter},
  journal={The American Journal of Human Genetics},
  volume={73},
  number={5},
  pages={1162--1169},
  year={2003},
  publisher={Elsevier}
}

@article{zheng2015reconstruction,
  title={Reconstruction of genome ancestry blocks in multiparental populations},
  author={Zheng, Chaozhi and Boer, Martin P and van Eeuwijk, Fred A},
  journal={Genetics},
  volume={200},
  number={4},
  pages={1073--1087},
  year={2015},
  publisher={Genetics Soc America}
}

@article{marchini2006comparison,
  title={A comparison of phasing algorithms for trios and unrelated individuals},
  author={Marchini, Jonathan and Cutler, David and Patterson, Nick and Stephens, Matthew and Eskin, Eleazar and Halperin, Eran and Lin, Shin and Qin, Zhaohui S and Munro, Heather M and Abecasis, Gon{\c{c}}alo R and others},
  journal={The American Journal of Human Genetics},
  volume={78},
  number={3},
  pages={437--450},
  year={2006},
  publisher={Elsevier}
}

@article{abecasis2002merlin,
  title={Merlin—rapid analysis of dense genetic maps using sparse gene flow trees},
  author={Abecasis, Gon{\c{c}}alo R and Cherny, Stacey S and Cookson, William O and Cardon, Lon R},
  journal={Nature genetics},
  volume={30},
  number={1},
  pages={97--101},
  year={2002},
  publisher={Nature Publishing Group}
}

@article{aylor2011genetic,
  title={Genetic analysis of complex traits in the emerging Collaborative Cross},
  author={Aylor, David L and Valdar, William and Foulds-Mathes, Wendy and Buus, Ryan J and Verdugo, Ricardo A and Baric, Ralph S and Ferris, Martin T and Frelinger, Jeff A and Heise, Mark and Frieman, Matt B and others},
  journal={Genome research},
  volume={21},
  number={8},
  pages={1213--1222},
  year={2011},
  publisher={Cold Spring Harbor Lab}
}

@article{broman2019r,
  title={R/qtl2: software for mapping quantitative trait loci with high-dimensional data and multiparent populations},
  author={Broman, Karl W and Gatti, Daniel M and Simecek, Petr and Furlotte, Nicholas A and Prins, Pjotr and Sen, {\'S}aunak and Yandell, Brian S and Churchill, Gary A},
  journal={Genetics},
  volume={211},
  number={2},
  pages={495--502},
  year={2019},
  publisher={Genetics Soc America}
}

@article{churchill2004collaborative,
  title={The Collaborative Cross, a community resource for the genetic analysis of complex traits},
  author={Churchill, Gary A and Airey, David C and Allayee, Hooman and Angel, Joe M and Attie, Alan D and Beatty, Jackson and Beavis, William D and Belknap, John K and Bennett, Beth and Berrettini, Wade and others},
  journal={Nature genetics},
  volume={36},
  number={11},
  pages={1133},
  year={2004},
  publisher={Nature Publishing Group}
}

@article{davies2016rapid,
  title={Rapid genotype imputation from sequence without reference panels},
  author={Davies, Robert W and Flint, Jonathan and Myers, Simon and Mott, Richard},
  journal={Nature genetics},
  volume={48},
  number={8},
  pages={965},
  year={2016},
  publisher={Nature Publishing Group}
}

@article{ganal2011large,
  title={A large maize (\textit{Zea mays} L.) SNP genotyping array: development and germplasm genotyping, and genetic mapping to compare with the {B73} reference genome},
  author={Ganal, Martin W and Durstewitz, Gregor and Polley, Andreas and B{\'e}rard, Aur{\'e}lie and Buckler, Edward S and Charcosset, Alain and Clarke, Joseph D and Graner, Eva-Maria and Hansen, Mark and Joets, Johann and others},
  journal={PloS one},
  volume={6},
  number={12},
  year={2011},
  publisher={Public Library of Science}
}
