## Supplementary figures and images for "SPEARS: Standard Performance Evaluation of Ancestral Reconstruction through Simulation"

### Figure_1.pdf

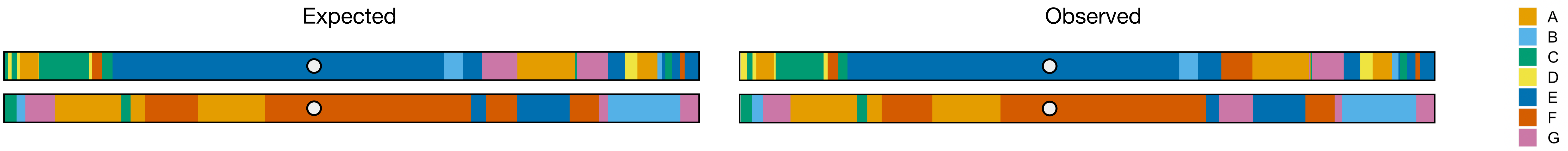

### Figure_1.png

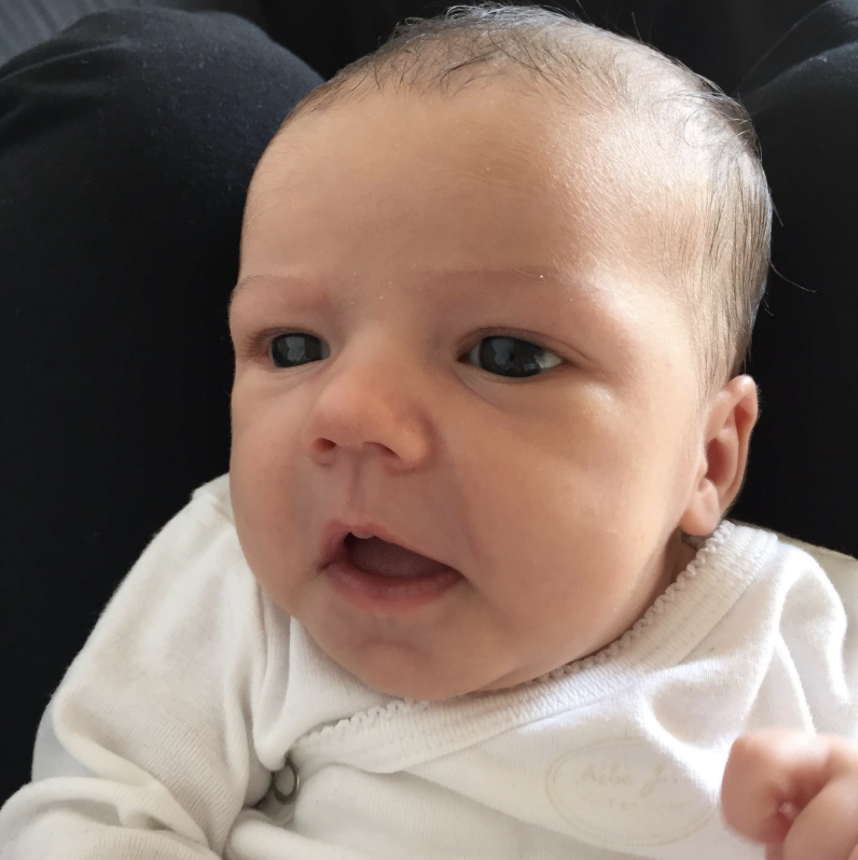
